# A Functional Basis for the Developmental Sequence of the Macrostructure of the Venus Flower Basket (*Euplectella aspergillum*)

**DOI:** 10.64898/2026.02.05.704042

**Authors:** Y. Mistry, S. Morankar, D. Kingsbury, N. Chawla, C. A. Penick, D. Bhate

## Abstract

Despite receiving significant interest from the biological and engineering communities, several questions about the underlying reasons for the form of the deep-sea sponge Venus Flower Basket (*Euplectela aspergillum)* remain unanswered. In particular, the basis for the sequence of emergence of three distinct macroscopic geometric features, while speculated upon, has not been validated. These features are (i) an interwoven cross-grid in the juvenile stage, (ii) a diagonal weave atop this initial grid, and (iii) a helical ridge that emerges in the mature phase of the sponge. This work uses computational design and additive manufacturing to fabricate models of each of these phases in sequence and subjects the models to mechanical tests in compression, bending and torsion. The results show that each feature has a singular advantage, even after accounting for the increase in mass associated with their addition: increased compliance for interwoven cross-grid, increased bending stiffness for diagonal weaves, and improved torsional rigidity for the helical ridge. The work postulates that the prioritization of compliance in the juvenile phase and a transition to a stiffer structure in the mature phase is a strategy that enables the sponge to avoid high internal stresses to avoid failure in its inherently brittle, silica-derived architecture.

**Highlights:** 1. The benefits of three macrostructural design elements of the Venus Flower Basket (*E. aspergillum*) that emerge sequentially in its development are examined: (i) the interweaving cross-grid, (ii) the diagonal weave that overlays this cross-grid, and (iii) the helical ridge reinforcement that forms over this diagonal weave.

2. For the first time, this work models each of these design features sequentially and studies their mechanical benefits through an experimental exploration of the behavior of idealized geometries in three test domains: compression, bending, and torsion.

3. Results show that each of these three structural elements has a unique functional advantage depending on the organism’s growth stage, even after accounting for differences in mass: interweaving enables higher compliance under compression, the diagonal weave improves bending stiffness, and the helical ridge improves torsional stiffness.

## 1. Introduction

The structure of life has fascinated not just biologists, but also mathematicians, physicists, and engineers [1]. This fascination beyond the domain of its immediate study is in part attributable to the beauty and complexity of natural form, and its remarkable ability to meet multiple functional objectives. Identifying structure-function relationships remains a key aspect of the biological sciences [2]. Establishing these relationships is also a fundamental requirement for abstracting design principles for bio-inspired design [3]. However, the very basis of fascination - the geometric and functional complexity of biological form, also poses a challenge when attempting to establish a functional basis for its evolution. The focus of this work - *Euplectella aspergillum*, commonly known as the Venus Flower Basket, is one such example. *E. aspergillum* is a deep-sea glass sponge from the class *Hexactinellida*. It has received significant interest from the scientific and engineering communities over the past two decades [4], much of it precisely due to its complex and hierarchical structure [5]. Despite being the subject of numerous studies (tabulated in **Supplementary Material - Table S1**), central questions about the underlying reasons for the growth and form of *E. aspergillum* remain unresolved.

*E. aspergillum* is typically found in the Pacific and Indian oceans, covered in a mat of soft, filter-feeding tissue, growing at depths of 100 - 1000 m where the temperature is between 2-11°C [6] [7]. At these depths, the structure experiences lateral forces from water currents and is at risk of predation by vertebrates, while managing forces of swimming fish making contact with them [8]. Given the inherently brittle nature of the silica-based material that the basket is comprised of [4] [9] and their remarkably long lives —estimated at over 11,000 years for some relatives [10]—it is remarkable that this structure has thrived in these conditions. It has been hypothesized that this exceptional behavior is enabled by hierarchical design strategies spanning length scales from nanometers to centimeters [11]. At the nanoscale, the growth of *E. aspergillum* starts with the deposition of silica particles measuring 3 nm in diameter around proteinaceous axial filaments [12], which form concentric cylinders of 50-200 nm in diameter [13] [14]. This formation leads to the development of struts with thicknesses of 80-100 µm [5]. The cross-section of these struts consists of a concentric, laminated core which is critical for imparting damage tolerance to individual spicules and the overall skeletal structure [5] [15] [16]. The spicules grow in length [17] to form a series of overlapping vertical, horizontal, and diagonal struts on a uniform cross-grid mesh averaging 2.5–3 mm in mesh size [5].

In addition to these nano, micro and meso-scale features, the mature *E. aspergillu*m also reveals three macrostructural design features shown in Figure 1a: (i) an interweaving cross-grid, (ii) a diagonal weave that is intertwined with the cross-grid, and (iii) a helical ridge that forms over the diagonal weave. Several studies have examined the structural benefits of these three design features, but with limiting assumptions. For example, studies have shown that the cross-grid and diagonal weave improve mechanical properties of the structure [18] [19] [20] [21], but these have been conducted on two-dimensional extruded structures which do not mimic the mechanics associated with the cylindrical nature of the macrostructure. Further, most prior studies also ignore the fact that both the cross-grid and the diagonal weave are not fully coupled at every node and have a weave-like quality [8].

**Figure 1.**
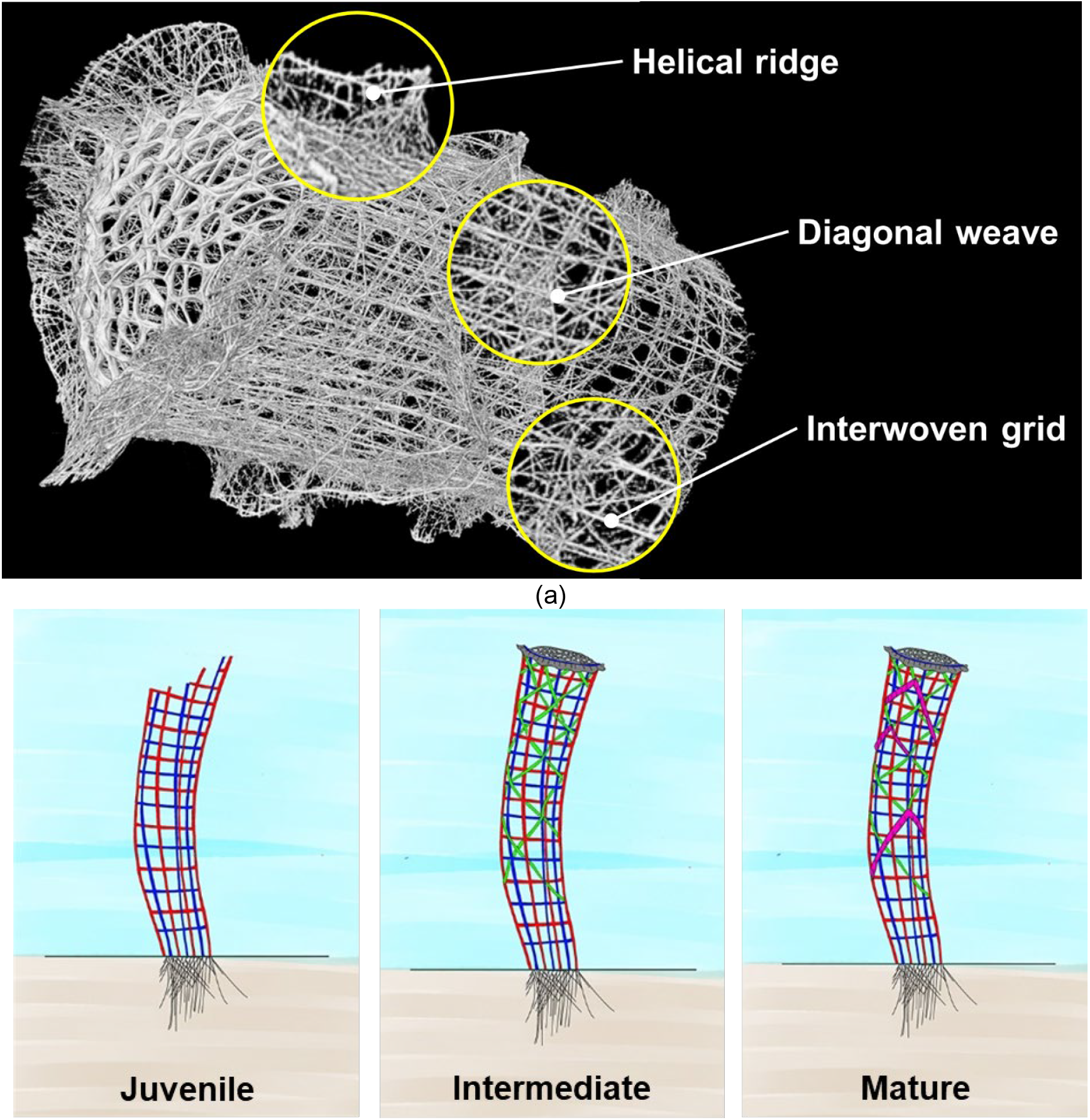
**(a)** *E. aspergillum* skeleton structure, indicating the three structural elements that are the focus of this work: the interwoven grid, the diagonal weave and the helical ridge; and **(b)** the three phases of growth at a macrostructural level of the *E. aspergillum*

An interesting aspect of the macrostructure of the *E. aspergillum*, which has received little prior study, is the sequence in which each of these three features appears. The long, multi-century life spans of this organism, and the difficulty of deep-sea field studies has made it challenging to record the sequence of growth. However, rational arguments and empirical observations in specimens suggest that the cross-grid forms first, followed by the diagonal weave, and eventually the helical ridges [22] (Figure 1b). This is reasonable, given the physical layering of the latter structure over the former, as visible in specimens. What is still unresolved is the functional basis for this particular sequence of development. For example, studies have identified and discussed the benefit of the helical ridge, which has been shown to improve torsional rigidity and provide hydrodynamic benefits to the structure [5] [22] [19] [23]. But why does the helical ridge only develop for mature *E. aspergillum* – how is the organism able to survive without a ridge in its juvenile state? It is questions of this nature that this work aims to answer by examining the mechanical behavior of designed structures that embody each of these three features as design principles and testing hypotheses for their developmental sequence.

The three hypotheses this work seeks to test are as follows:

i. The interwoven grid forms at the early stage of growth in the juvenile *E. aspergillum* to give it compliance under deformation while the structure is not fully developed.
ii. The diagonal weave develops next as *E. aspergillum* grows taller, to protect the structure against bending stresses that grow proportionally with length.
iii. The helical ridge emerges at a later stage as *E. aspegillum* grows in diameter (girth), to protect the structure against torsional loads, the stresses from which grow in proportion to diameter.

Crucially, and in differing from prior work, the studies here are conducted (i) on cylindrical specimens that better approximate the geometry of *E. aspergillum*; (ii) on geometries where features are added in the same sequence of growth as shown in Figure 1b; and (iii) the decoupled, interwoven nature of the lattice grid is represented as such and not projected onto a plane. This work represents, to the best of the authors’ knowledge, the first study that demonstrates the structural basis for the sequence in which the macrostructural features in *E. aspergillum* appear. Section 2 details the methods used to conduct this study. Section 3 compiles all the test results from the study and is followed by Section 4 which considers the validity of the three hypotheses identified in this work. Section 5 concludes this work with a summary of the key findings and their implications for future research.

## 2. Methods

The primary objective of this work is to test the three hypotheses listed previously, which essentially relate structure to function, but in the context of the organism’s development. Lauder proposed that a structure-function relationship in biology can be validated in two ways: using a phylogenetic comparative method, or a paradigm method [24]. In the former, more common when applied to extinct species, emphasis is placed on identifying homologous structures in related species and leveraging known relationships in them to establish confidence in the prediction of a similar function for the structure and species of interest. The paradigm approach involves developing abstractions or models that can be used to make structure-function predictions, and comparing these predictions of structure to the one observed in reality [25]. These models can either be analytical, computational or, as in the approach taken in this work, involve an experimental study of a representative 3D model as a basis for testing hypothesized relationships between structure and function. The approach in this work is thus a milder form of the paradigm method – instead of making predictions of structure given a functional objective and constraints, it merely seeks to validate a hypothesized relationship between structure and function by testing a structure of interest. While not providing the same level of confidence as the paradigm method, it nonetheless is a useful addition to the accumulation of evidence in support of a particular structure-function relationship, as will be argued in this work for *E. aspergillum*.

This section briefly describes the main methods employed to arrive at experimental results, beginning with X-ray tomography of the *E. aspergillum*, followed by the representation of its key design features in a 3D model, manufacturing using a 3D printing process, and finally, experimental testing in different mechanical loading conditions.

### 2.1 X-Ray Tomography

The structure of E. *aspergillum* was first investigated with a Zeiss Xradia Versa 620 x-ray scanning microscope, using a specimen obtained online. Several scans with pixel size ranging from 23.5 µm to 8.4 µm were performed to investigate the structure at a range of length scales. After scanning, all 3D datasets were processed in an image analysis software called Avizo 9.0 for visualization and further analysis. More details on specimen sourcing, tomography setup and data analysis, as well as the findings from this part of the work have been published elsewhere [4] and are as a result not additionally addressed in this paper.

### 2.2 Design

All specimens in this study were designed using Grasshopper, which is a visual programming language and environment that runs within the Rhinoceros 3D computer-aided design (CAD) application and is commonly used by architects, jewelry designers and artists [26]. As shown in the Grasshopper script layout in Figure 2 for one instance of specimens developed in this work, design instructions can be grouped into component blocks, the outputs of which serve as inputs of other components. In this work, a range of primary input parameters are defined by the user, which are then passed downstream to modules that then generate the desired designs (**Grasshopper scripts used in this work are available as supplementary material**).

**Figure 2.**
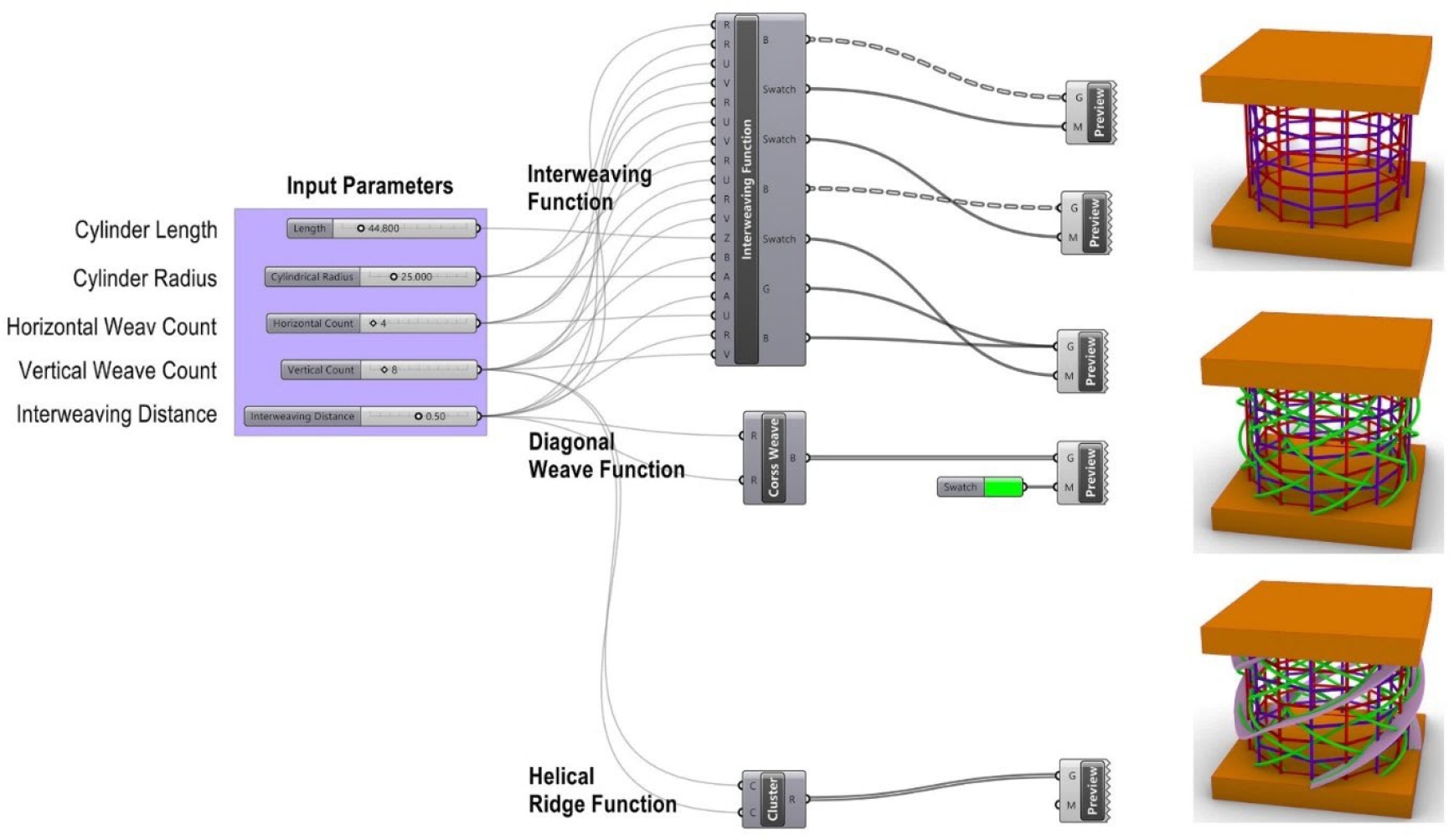
The Grasshopper script used in this work and examples of three types of specimens formed using the script – shown here for compression testing

### 2.3 Manufacturing

Selective Laser Sintering (SLS) using an EOS Formiga P110 machine, was used to manufacture specimens due to its ability to create complex structure without needing supports [27]. The material used was Polyamide 12 (PA12) [28]. Standard, supplier provided parameters were used to manufacture the part, and all standard post-processing steps were followed for optimum powder removal (steps shown in **Supplementary Figure S1**).

### 2.4 Mechanical Testing

The specimens were subjected to three different loading conditions: compression, 4-point bend, and torsion tests. These three loading conditions were selected to enable evaluation of specific hypotheses that are discussed in the next section. Compression tests were performed with a compression rate of 1mm/min, and the specimens were compressed till densification. The four-point bend test was performed based on ASTM D6272 [29] with a 1mm/min loading rate. A video extensometer reading off a mechanical plunger was used to measure the flexural displacement. Load was applied until the flexural displacement limit reached the mechanical plunger limit of 15mm. Compression and four-point bending tests were conducted on an Instron 5985 electromechanical universal testing machine. Torsion tests were performed guided by ASTM E143 [30] with a 0.3 degree per second testing rate, with specimens twisted to 180 degrees, and free to move in the axial direction to limit deformation mode to torsion. The torsion test was performed on a 311 series frame Universal Testing Machine (UTM) manufactured by TestResources, Inc. The UTM had a torsion add-on with a 2000 lb axial load and 1128 in-lb torsional load maximum.

## 3. Design Principle Abstraction

A key aspect in the methodology of bio-inspired design is the abstraction of a design principle that enables translation of a structure-function relationship in biological form to an application of interest in engineering [31]. However, this approach can also be used when the underlying biological structure-function relationship is not well established. Common to both is the need to abstract a design feature that can be represented using engineering design tools. For this work, x-ray microtomography was used to abstract three design features in *E. aspergillum* and enable subsequent study. The results from the abstraction are discussed in this section for each of the three design features in turn.

### 3.1 Interwoven Cross Grid (Juvenile Phase)

A close examination of the lattice-like grid of the *E. aspergillum* reveals that it is not a fully connected lattice but consists instead of two lattices interwoven with each other as shown in Figure 3a, consistent with observations in the literature [5]. This design feature was implemented in Grasshopper as shown in Figure 3b and fabricated with Selective Laser Sintering (Figure 3c) such that there was sufficient clearance to ensure there was no sintering of decoupled beams to each other. This follows prior work where this concept was first explored for 3D lattices [8].

**Figure 3.**
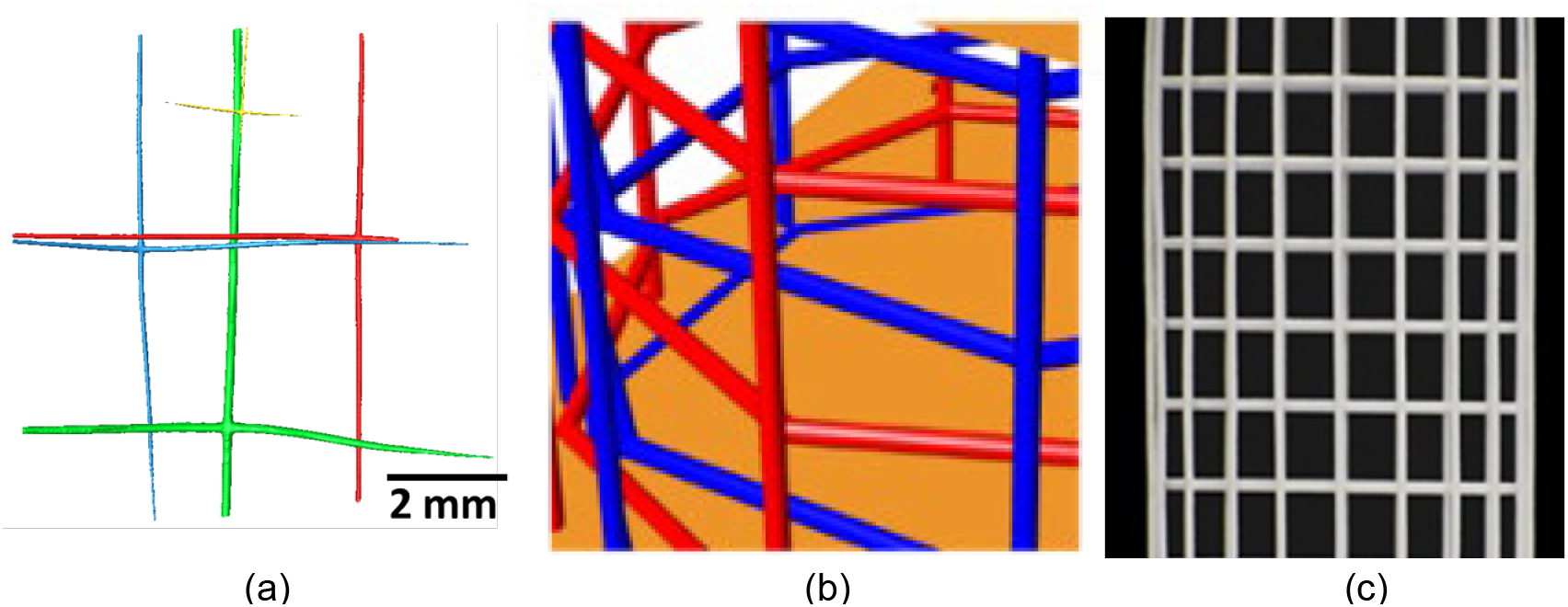
X-ray microtomography image showing the lattice-like cross grid of the *E. aspergillum*; (b) abstraction of this design feature in Grasshopper; and (c) 3D-printed specimen with interweaving cross struts

### 3.2 Diagonal Weave (Intermediate Phase)

The diagonal weave observed in the Venus flower basket (Figure 4a) is interwoven with the cross grid discussed above. XCT data revealed no consistent and definite pattern in the weaving of diagonal struts. However, the majority of struts were observed to pass under one vertical strut after passing over two vertical struts [5]. This pattern was replicated in Grasshopper with cross-diagonal struts making contact every two passes as shown in Figure 4b, where the green struts indicate the diagonal weave, overlaid on top of the blue and red interwoven cross-grid. One of the specimens fabricated our of SLS using this design principle is shown in Figure 4c.

**Figure 4.**
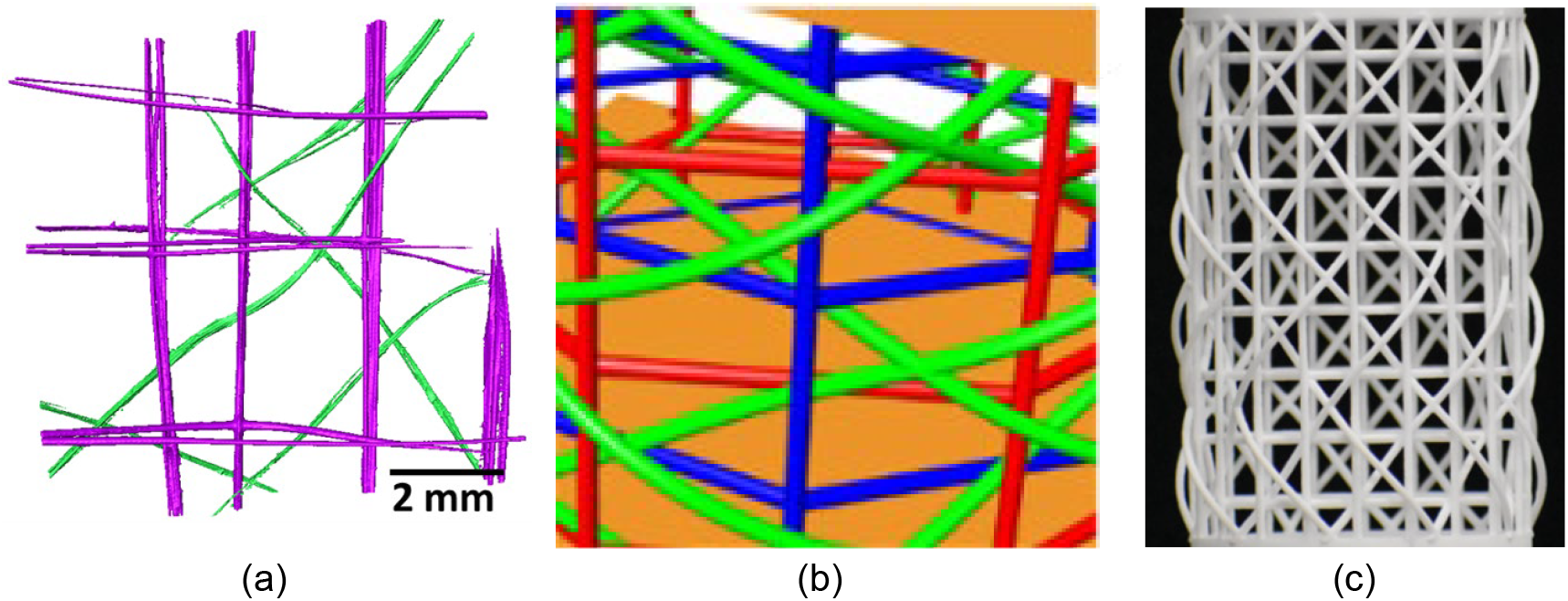
(a) X-ray microtomography image showing the diagonal struts interwoven with the cross grid of the E. aspergillum; (b) abstraction of this design feature in Grasshopper; and (c) 3D-printed specimen with interweaving cross- and diagonal members

### 3.3 Helical Ridge (Mature Phase)

The final design feature explored in this work is the helical ridge, which emerges perpendicular to the plane of the basket wall, as shown in Figure 5a. These ridges are formed along the diagonal strut, which itself forms helically around the cylindrical basket. The helical ridge also displays a particular pattern: it forms on top of every other diagonal weave and makes a 90 degree turn to loop around itself [19]. These details were idealized in the grasshopper design (Figure 5b) and realized with SLS 3D printing (Figure 5c).

**Figure 5.**
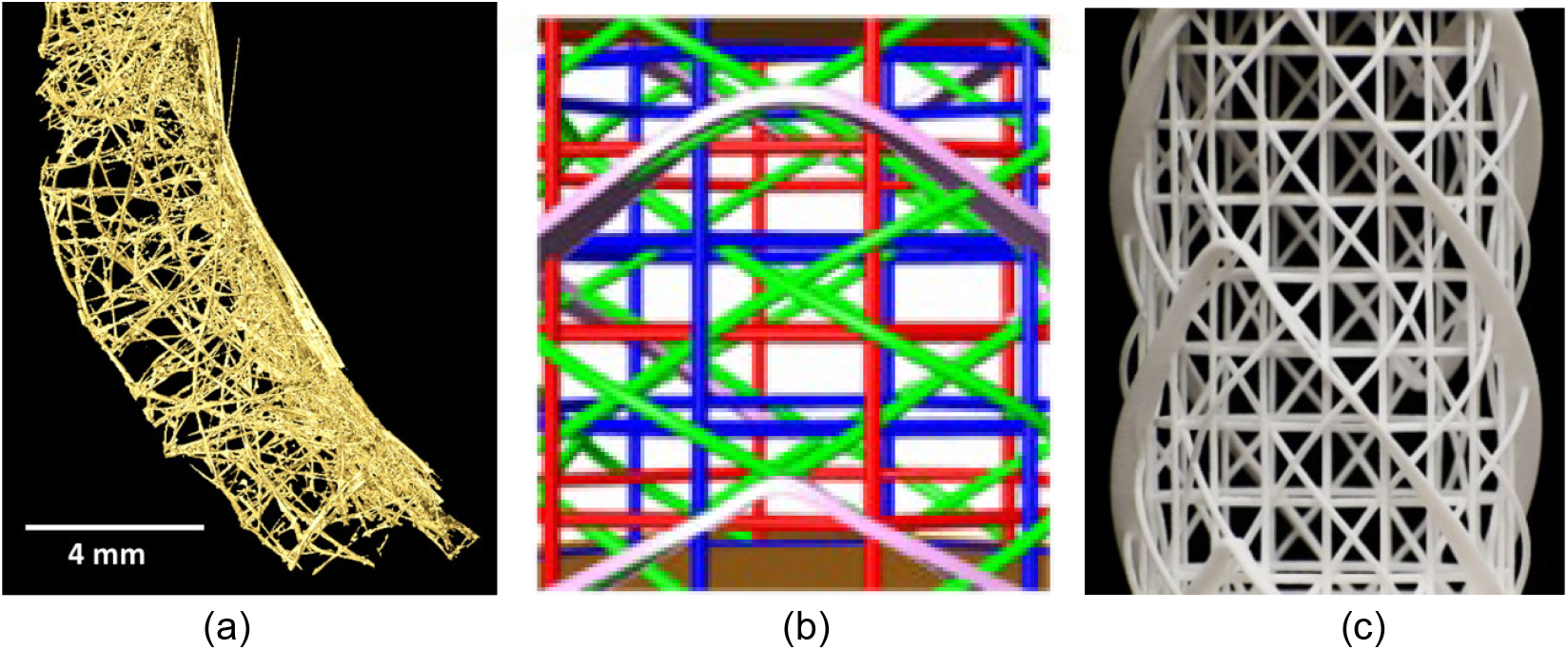
(a) X-ray microtomography image showing the helical ridges as the project normally outward from the cylindrical macrostructure; (b) abstraction of this design feature in Grasshopper; and (c) 3D-printed specimen with helical ridges atop interweaving cross- and diagonal members

With the specimens designed and fabricated, an experimental protocol was developed to test the afore mentioned hypotheses, for which three test conditions were identified: compression, bending, and torsion. These three conditions were selected based on the eco-mechanical environment [32] that *E. aspergillum* likely experiences during its development. It grows to a height of 10 to 30 cm and is only anchored to the ocean bed using a holdfast apparatus at one end [4], suggesting vulnerability to bending, particularly as it grows in length. It is also likely that the structure experiences torsional forces due to turbulence induced non-uniform loads, and predation from fishes and other marine life [28].

A total of 12 specimens were designed for this study: four types of geometries (baseline, interweaving, diagonal weave and helical ridge) for each of the three test conditions (compression, 4-point bending, and torsion), and are shown collectively in Figure 6, with the experimental setups for the 3D-printed specimens shown in Figures 7a-7c for each of the three test conditions. A total of 3 specimens (i.e. 2 replicates) per geometry and test condition were tested. The baseline design was designed as a fully connected lattice grid, with no weaving or ridges.

**Figure 6.**
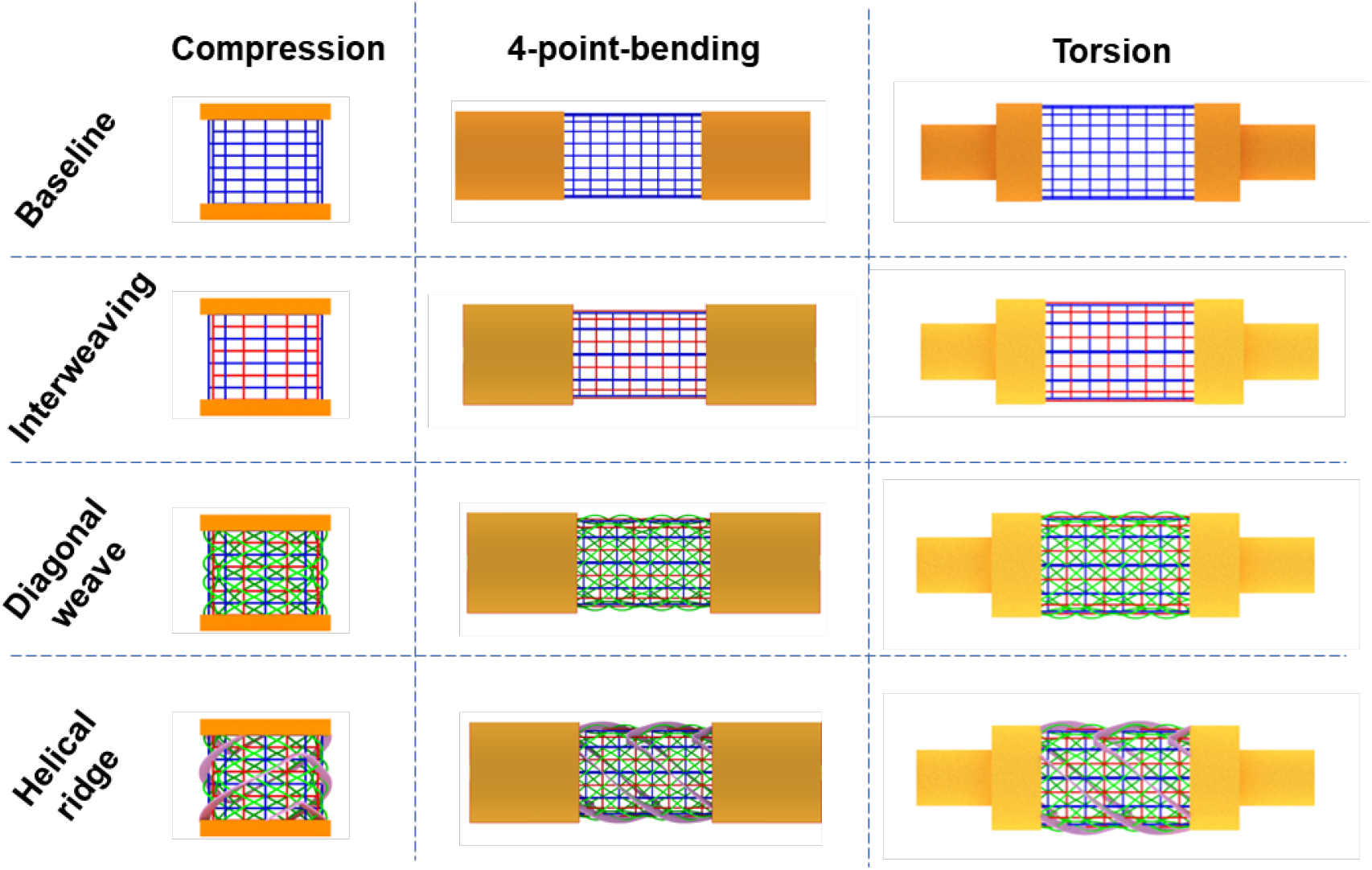
Grasshopper generated 3D design files for each of the three test conditions, across all four specimen designs

**Figure 7.**
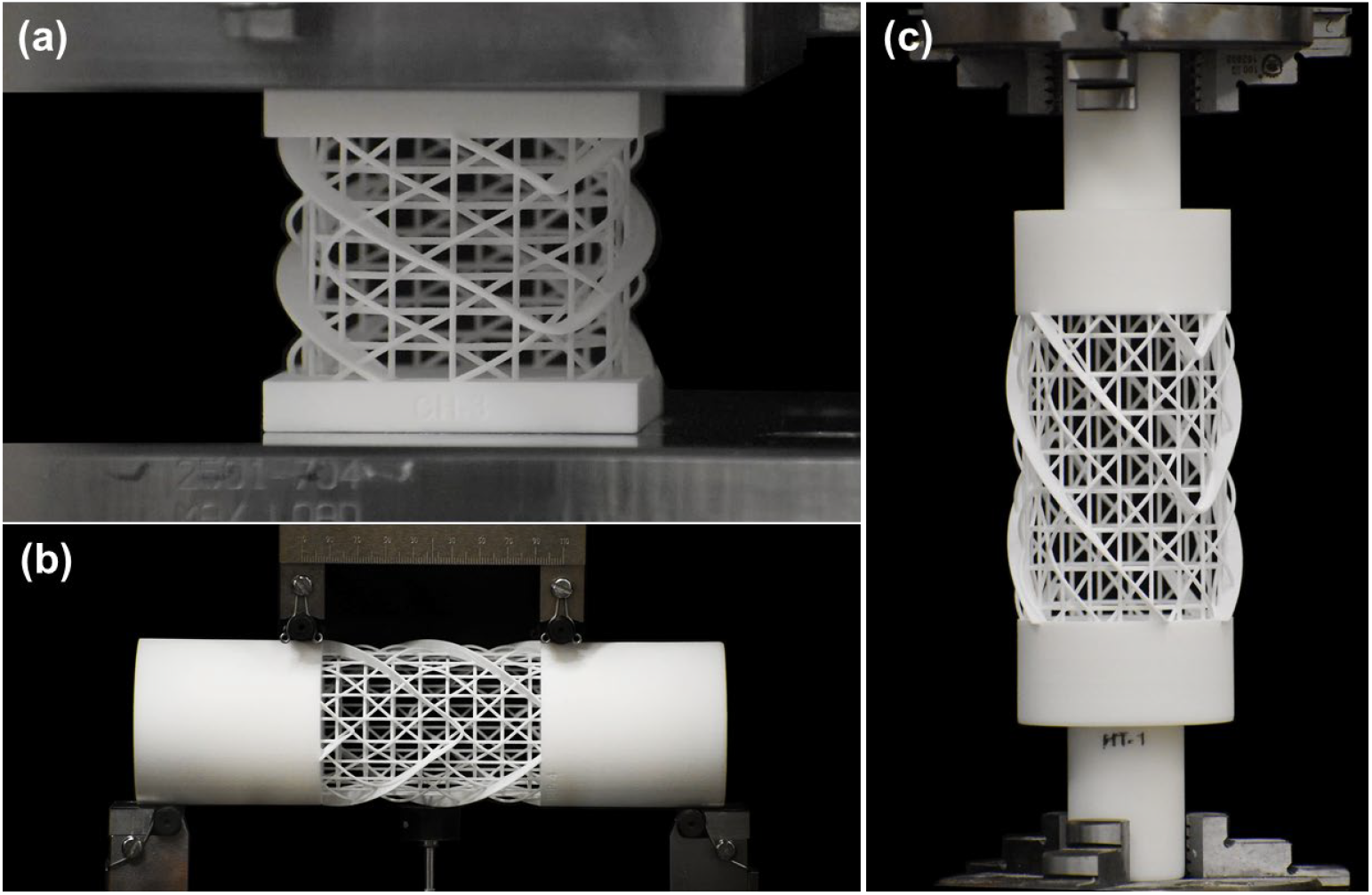
3D printed test specimen setup for the (a) compression test, (b) 4-point bend test, and (c) torsional test – shown here for the helical ridge specimens only

## 4. Results

The results from the compression, 4-point bend, and torsion tests are discussed in turn below, with special emphasis paid to whether any particular design performed better than the others for a given test condition. The following section analyzes these results in the context of the hypotheses presented at the start of the paper and discusses their implications for our understanding of the basis for the sequential growth of *E. aspergillum*.

### 3.2.1 Compression

Figure 8a shows the raw load-displacement data for all four specimen designs under compression loading, the spread representing the variation across the replicates. Among all designs evaluated, the interweaving specimens had the lowest peak load and demonstrated a stress plateau with the fewest undulations, properties desirable in energy absorption [33]. The lower peak load of the interweaving specimen is expected on account of the larger beam lengths resulting from the decoupled nodes, effectively resulting in increasing beam length [8]. Per Euler beam buckling theory, the force required to buckle a column is inversely proportional to the square of its length. Hence, the peak force for the fully connected baseline specimen with a shorter beam length is higher than the interweaving specimen.

**Figure 8.**
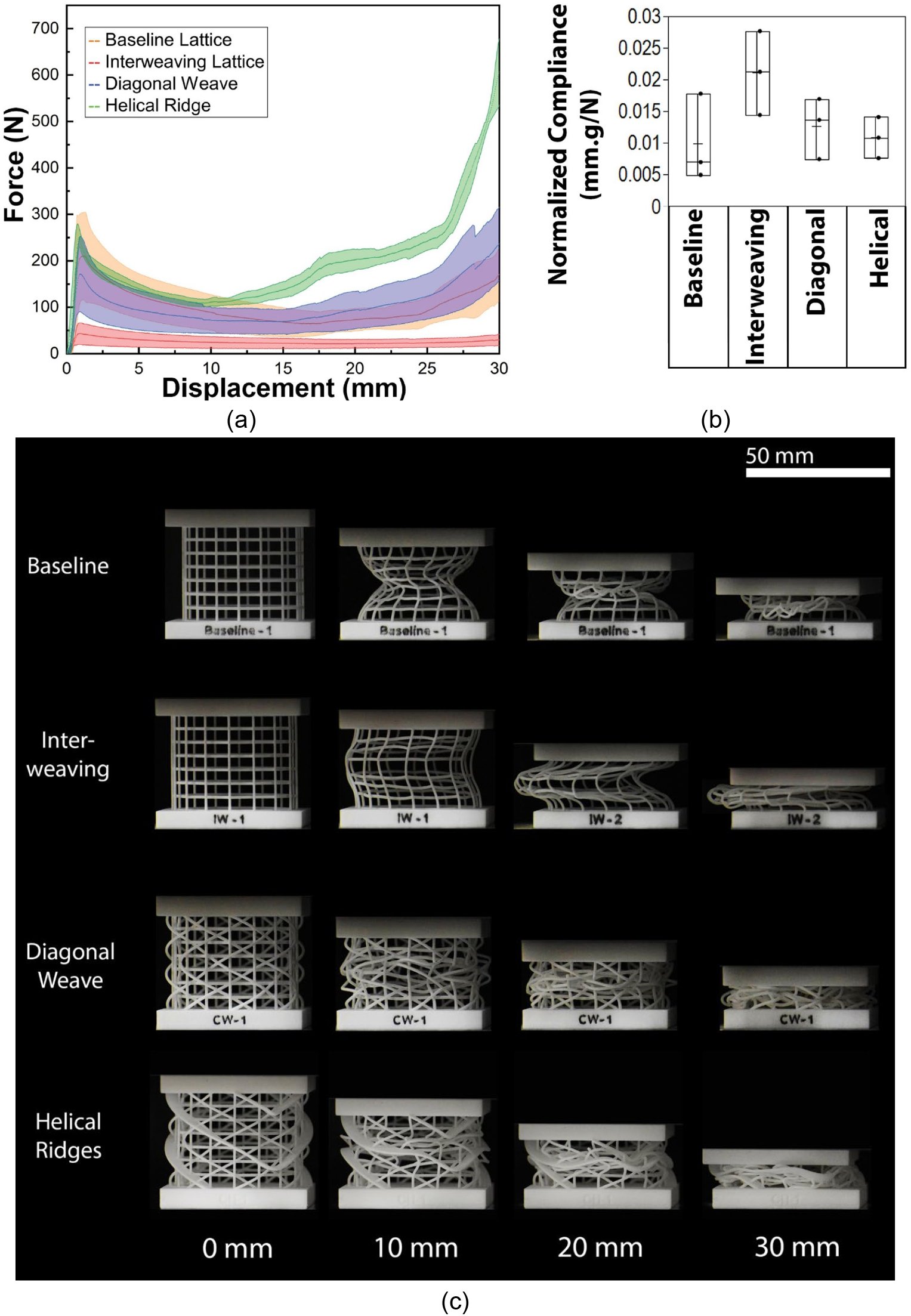
Compression test results for the four designs evaluated in this study: (a) force-displacement response, (b) normalized compliance for all four designs, showing highest values for the interweaving structure, and (c) deformation patterns of each specimen at four different levels of displacement

Of greater interest to this work, however, is the compliance of these structures under compression, in line with the first stated hypothesis associated with the juvenile state of the sea sponge. Compliance is the inverse of stiffness (stiffness, in turn, is the ratio of force over displacement), and is estimated in the initial, linear elastic response of the force-displacement graph. To enable meaningful comparisons across the four design types with different masses, stiffness is commonly normalized (divided) by the mass of the structure to estimate what is called a “specific stiffness”. an engineering perspective, where specific stiffness (per unit mass) is a key metric of interest – several applications such as aircraft components demand high stiffness with low mass. There are, however, certain applications where high compliance is desired, in energy absorbers and piezoelectric sensors, for example.

For the purposes of this work, the inverse of the stiffness/mass ratio was used to estimate a normalized compliance, which is shown in Figure 8b for all four designs. The comparison analysis of compliance shows that interweaving structures have a higher normalized compliance than the other structures. This is in line with the authors’ prior work showing a similar strategy implemented for 3D lattices inspired by *E. aspergillum*, that showed that creating interweaving lattices by decoupling nodes enabled significant increases in mass-normalized compliance [8]. Deformation patterns under compression for each of the four designs are shown in Figure 8c at four different levels of displacement. The baseline structure, with a fully connected grid, deforms with an hourglass-like form, while the diagonal weave and helical ridge maintain the external cylindrical shape while undergoing collapse. In contrast to these three shapes, the interweaving structure experiences significant buckling in its individual longer beams and demonstrates a buckling-like response in its macro-structure.

### 3.2.2 Bending

Bending is an important mode of loading for structures because it induces non-uniform stress distributions that can result in higher maximum stresses compared to those produced by the same magnitude of axial loads (tension or compression). Results obtained from 4-point bending tests conducted on the four different designs are shown in Figures 9a-9c. Figure 9a shows the flexural (bending) load-displacement data for all four designs obtained from the bending test, with the scatter on account of the differences in the three samples tested for each design.

**Figure 9.**
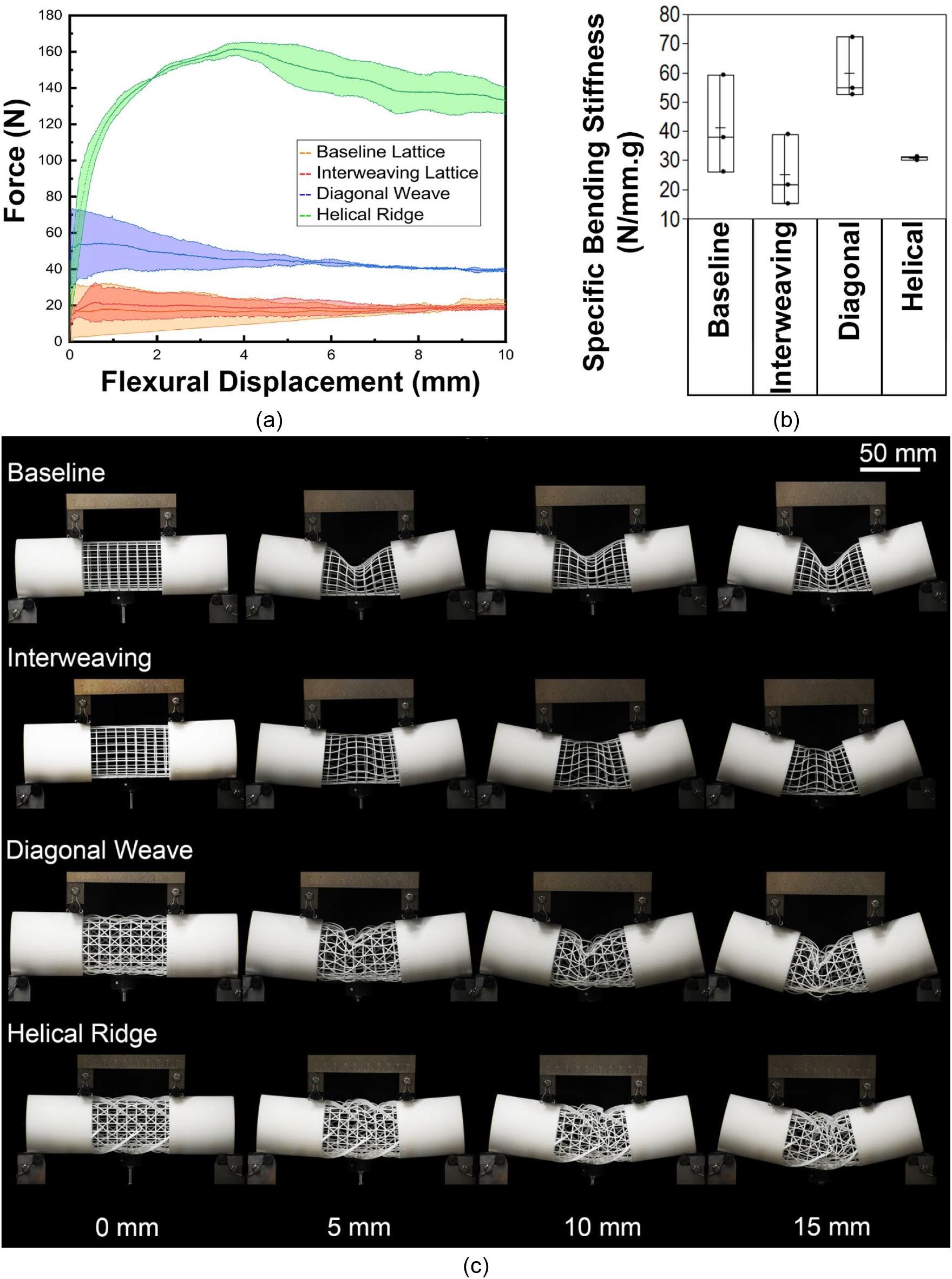
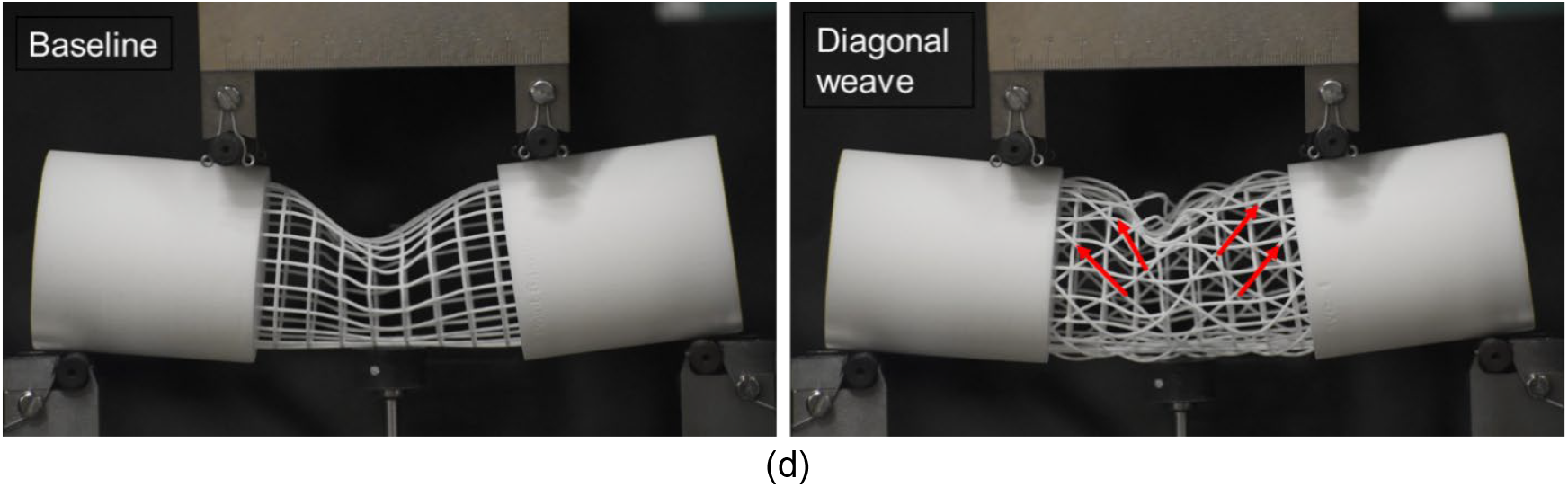
Four-point bend test result for four types of design specimens: (a) shows the force-displacement data, (b) shows mass-normalized bending stiffness, (c) deformation of specimens at different downward displacement values, and (d) increase stiffness of the diagonal weave under bending may be attributed to the presence of diagonal struts that are loaded axially

The data points to two notable observations: first, the helical ridge structure clearly shows the highest load bearing capacity, which holds true even after accounting for the additional mass. Secondly, of the four designs, the diagonal weave was found to be stiffest under bending in relation to the other designs, with the average mass-normalized specific bending stiffness being 62% stiffer than the baseline, as shown in Figure 9b. Deformation patterns under bending are shown in Figure 9c at different values of applied displacement. The baseline, interweaving and diagonal specimens all show an hour-glass-like buckling collapse like that seen in compression for the baseline specimen (Figure 8c) – this is clearly visible for the baseline and diagonal specimens. The addition of the helical ridges seems to have the effect of delaying the onset of this collapse and allowing the structure to bear a higher peak load.

The increased stiffness (per unit mass) of the diagonal weave structure is somewhat unexpected but is likely due to the presence of the diagonal struts that experience axial loading while the structure is globally under bending. These struts increase the resistance to deformation. While the helical ridge specimens also include the diagonal weave, the additional mass does not bring any benefit in stiffness, but does enable larger load bearing capabilities by resisting the onset of buckling in the macro-structure.

### 3.2.3 Torsion

While axial (tension or compression) and bending loads are common for most structures, torsion tends to be relevant when a structure is supported at one end and prone to twisting loads being applied at the other – these loads are similar to bending loads, but are applied off-axis and as such result in a twisting of the structure instead of bending, which in turn results in shear deformation within the structure. A morphology such as *E. aspergillum*, with its hollow cylindrical nature and end-anchoring is a prime candidate for these loads.

To study the effects of torsional loading, an angular deformation (twist) was applied to the specimens and the reaction torque measured. Figure 10a shows this measured torque vs angle of twist for three replicates for each of the four designs. The helical ridge design had the highest peak torque in comparison to the other designs. This remains true even after accounting for the additional mass. As before, the effective torsional rigidity can be estimated for these structures, which is a measure of the stiffness of the structure under torsion. This quantity can be then divided by the mass of the structure to obtain a specific torsional rigidity (i.e. per unit mass) – this quantity is shown in Figure 10b for each of the designs studied, and shows the remarkable finding that the helical ridge design has the highest value in comparison to the others, and is nearly two orders of magnitude higher than the baseline design. The diagonal design also has a significantly higher specific torsional rigidity compared to the baseline, with the interweaving structure falling below it. An interesting observation was also made in the deformation patterns shown in Figure 10c: all designs show a collapse of the cylindrical form well before the end-of-loading 180 degree rotation, corresponding with the drop in peak torque seen in Figure 10a, with the exception of the interweaving specimen, which only delays this collapse till the very end of the loading profile.

**Figure 10.**
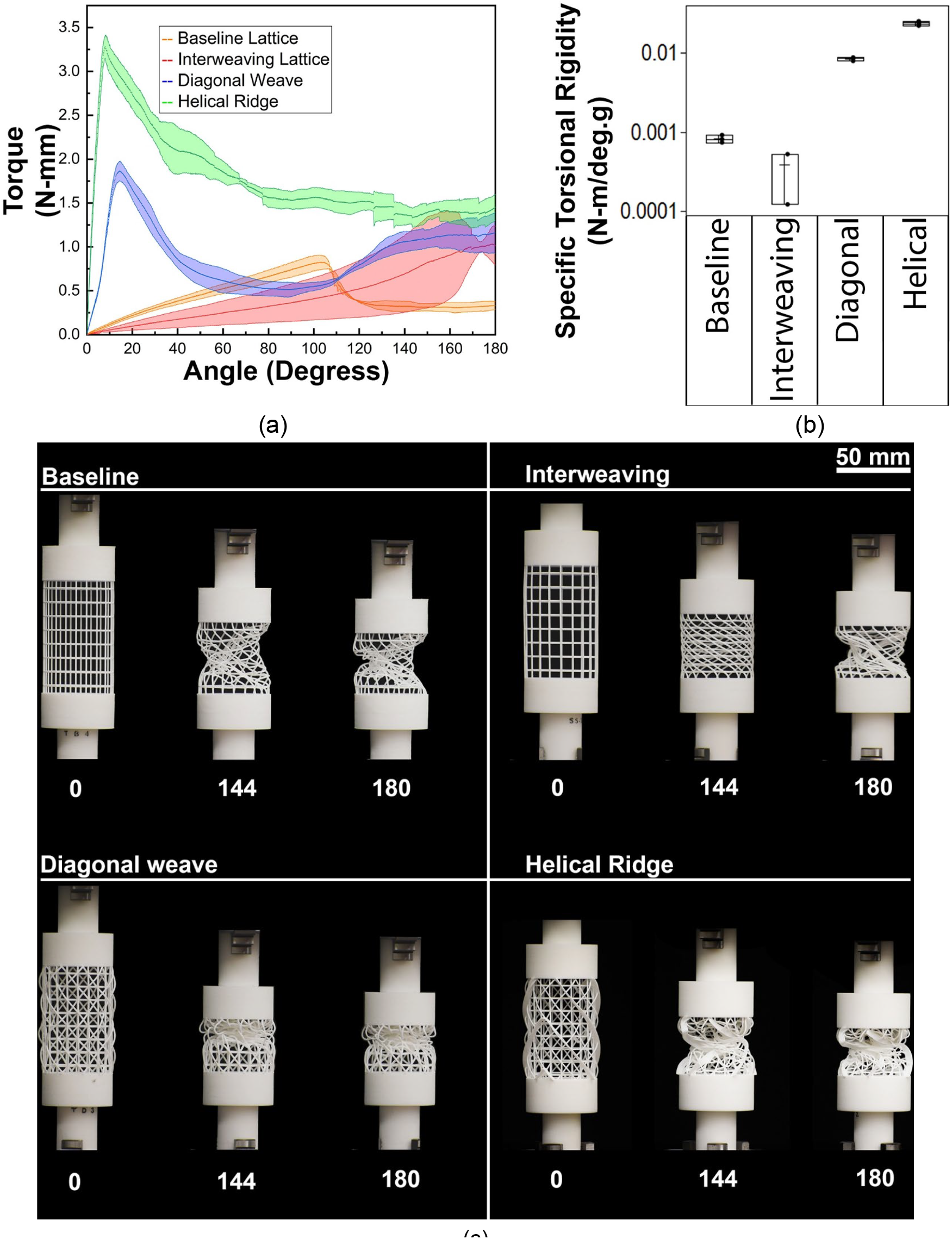
Torsional test results for the four types of designs, (a) torque vs twist angle, (b) specific torsional rigidity (log-scale), and (c) deformation patterns of specimens at different angles of twist

## 4. Discussion

While prior literature has concerned itself with establishing structure-function relationships for *E. aspergillum*, this work placed emphasis on addressing questions around the developmental sequence of three macrostructural features: the interweaving grid, the diagonal weave, and the helical ridge. Why do these structures emerge in the order that they do? Why does the organism not have all three features present at all stages of its development, instead delaying the onset of the latter features till a later time?

At the center of the hypotheses identified at the start of the work are two concepts: one rooted in materials science - strength, and the other in solid mechanics – stiffness, and their interplay with applied forces and resulting deformation. Sea sponges (phylum Porifera) secrete a variety of mineral skeletons made of calcite, aragonite, and/or silica [34]. The skeleton of the *E. aspergillum* is made of silica – an inherently brittle material – in other words, a material that fails suddenly when it reaches its failure stress (with little or no plastic, flow-like deformation prior to failure). Given this inherent limitation in the material, there are two macroscopic approaches to the design of a structure made of silica to reduce its susceptibility to failure: make it very compliant so that it deforms easily, or very stiff to increase its load carrying capacity while reducing its deformation under this load – in both cases, the objective is to keep localized stresses below the failure stress. An analog of the former strategy is a thin glass fiber, and of the latter is a solid block of glass.

The argument this work makes is therefore this: *E. aspergillum* starts its life with a skeletal structure that is highly compliant – it has less material to work with in its juvenile phase, and a rigid, fully interconnected grid at this stage would result in a structure that would generate high localized internal stresses that would result in crack formation and subsequent failure of the structure. This work predicts that the sponge therefore develops an interwoven structure instead, that increases compliance and avoids formation of these localized stresses. As the structure grows longer however, it increasingly becomes vulnerable to bending. To address this increase in bending stresses, a diagonal weave is generated – which still keeps the structure relatively compliant under compression, while increasing bending stiffness. As the structure grows further, it is capped at the top and increases in diameter as well, becoming increasingly vulnerable to torsional loads, which were less relevant when the grid was only loosely connected. The helical ridge now grows on top of the diagonal weave – this has the effect of greatly reducing the overall compliance of the structure but is achieved with the benefit of greatly increasing the stiffness under torsional loads.

While the data presented thus far emphasizes normalization by mass in order to estimate an effectiveness of the geometry independent of the amount of material utilized, another way to normalize the data is in relation to the baseline design with a fully interconnected cross-grid, to obtain a non-dimensional relative stiffness measure. This is shown in Figure 11. The results clearly back up the previously stated arguments: the interweaving grid has a stiffness lower than that of the baseline under compression, the diagonal weave increases relative stiffness under bending, and the helical ridge maximizes the relative torsional stiffness. These findings are even more remarkable given that the Y-axis representing relative stiffness is plotted as a log-plot. Finally, to bring all these findings together in one place, the relationships between the developmental phase of the *E. aspergillum* and the associated structural features and their hypothesized benefits is summarized in Table 1.

**Table 1.** Summary of hypothesized structure-function relationships by developmental phase of *E. Aspergillum* and the supporting evidence in favor of each hypothesis.

|  | Developmental Phase of *E. Aspergillum* |  |  |
| --- | --- | --- | --- |
|  | Juvenile | Intermediate | Mature |
| **Emergent Structural Feature** | Interweaving grid | Diagonal weave | Helical ridge |
| **Associated Functional Hypothesis** | Increases compliance while structure is developing | Protects structure against bending stresses which increase with length | Protects structure against torsional loads which increase with diameter |
| **Supporting Experimental Evidence** | Interweaving grid has the highest normalized compliance, over twice that of the fully connected grid | Diagonal weave has the highest specific bending stiffness, over twice that of the interweaving grid alone | Helical ridge has the highest specific torsional rigidity, over two orders of magnitude higher than the interweaving grid alone |

**Figure 11.**
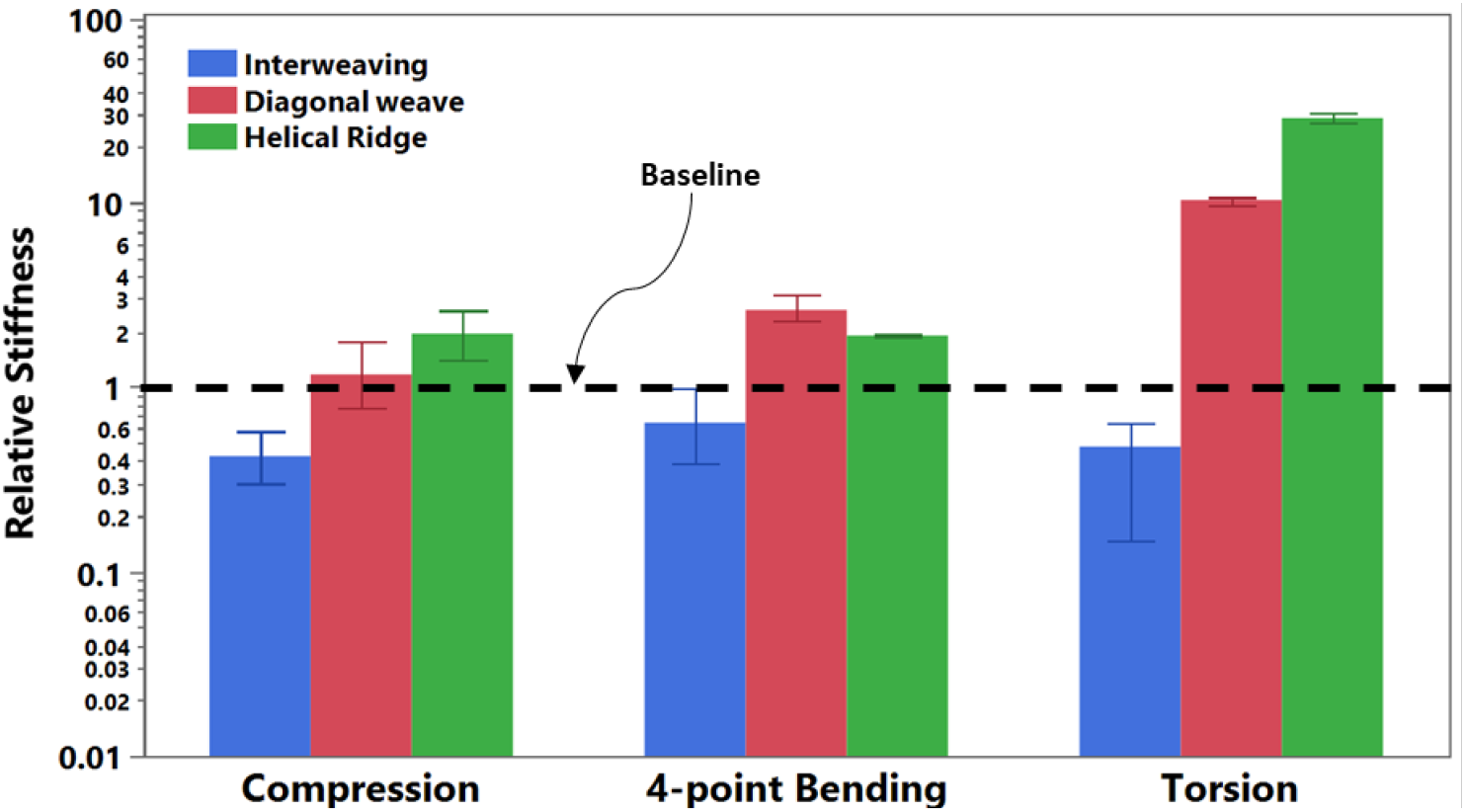
Normalized stiffness for each of the design features studied here as a function of the baseline (fully connected cross grid), for the three types of loading conditions studied in this work

## 4. Conclusions

This work attempted to establish a functional basis for the developmental sequence of the Venus flower basket (*E. aspergillum*), and also builds on prior work that specifically looked at one of the geometric features in more detail [8]. Two aspects of this collective work are worthy of summary: (i) the first has to do with the approach used, which has general applicability beyond the organism studied here, and (ii) the results obtained from the application of this approach to the development of *E. aspergillum*. With regard to the first aspect of the work, the following conclusions may be drawn:

- When attempting to attribute function to geometrically complex structures, one may first identify isolated forms of interest and then build models that sequentially layer them in turn, identifying the contribution of each form in isolation, as well as when combined with the other forms.
- Computational design tools can be used to generate these forms in a manner most relevant to the original organism and its environment, and then these generated designs can be used as the basis for either computational or experimental studies.
- Hypotheses relating structure to function can be first established knowing the eco-mechanical environment of the organism and verified using the developed models.
- These models can then be the basis for further prediction, and/or be extended beyond the domain of the organism into enabling engineering applications.

Specific to *E. aspergillum*, this work draws the following conclusions:

- The macrostructure development of *E. aspergillum* may be said to consist of three main phases, with a key feature emerging at each phase: the juvenile phase primarily consists of an interwoven grid, followed by the development of a diagonal weave at the intermediate phase, and culminating in a helical ridge at maturity.
- The interweaving grid enables high compliance under compression, minimizing local stresses at a time when the structure is relatively fine.
- The increase in length of the structure makes it vulnerable to bending-induced stresses, which are countered by the addition of a diagonal weave.
- Finally, the increase in girth and capping of the structure makes it more vulnerable to torsion-induced stresses, which are managed with the addition of a helical ridge.
- Remarkably, these enhancements in performance hold even after accounting for the additional mass associated with the forms needed to achieve them.

The approach developed in this work is not the only way to address challenges of unraveling geometric complexity in nature, nor are the structural benefits attributed to the geometric features a complete description of the basis for their evolution as it ignores hydrodynamic, biological and growth-related aspects. Nonetheless, it is hoped that this work is a useful addition to the growing literature on the truly full-of-wonders organism that is *E. aspergillum*.

## 6. Acknowledgements

This work was partially funded by the National Aeronautics and Space Administration (NASA) STTR program under contract 80NSSC18P2131 in support of the PeTaL (Periodic Table of Life) project. Dr. Vikram Shyam was the Principal Investigator for PeTaL and the Technical Monitor for the STTR phase 1 contract. Ezra McNichols was the Technical Monitor for the STTR phase 2 contract. The authors also acknowledge Dr. Cameron Noe for his efforts fabricating the structures used in this study with the Selective Laser Sintering process, using equipment at Arizona State University.

