## Supplementary table and figure for "A Functional Basis for the Developmental Sequence of the Macrostructure of the Venus Flower Basket (*Euplectella aspergillum*)"

**Contents:**

- Table S1
- Figure S1

**Table S1**. Summary of literature identifying and attributing elements

| Reference | Structural Element Described | Functional Basis Hypothesized | Validation |
|---|---|---|---|
| Morankar et al. (2022)<br>Sundar et al. (2003)<br>Fernandes et al. (2021)<br>Monn et al. (2015)<br>Chai et al. (2002)<br>Sarikaya et al. (2001)<br>Vangelatos et al. (2022)<br>Weaver et al. (2006) | Cylindrical Laminates/ End Spicules (Hold Fast Apparatus) | 1. Impart damage tolerance to the individual spicules.<br>2. Crack arrestation.<br>3. Energy dissipation.<br>4. Biological Optical Fibers | 1. In-Situ fiber tensile testing.<br>2. Simulation and numerical analysis.<br>3. Nano indentation, 3-point bend test on fibers (struts) |
| Mistry et al. (2023)<br>Weaver et al. (2006)<br>Brown et al. (2019)<br>Morankar et al. (2022)<br>Gray et al. (1867) | Interwoven lattice struts | 1. Increased compliance<br>2. Helps in stress distribution. | 1. Demonstrated, but in different geometry<br>2. Observed not yet validated. |
| Li et al. (2021)<br>He et al. (2021)<br>Weaver et al. (2006)<br>Fernandes et al. (2021)<br>Gray et al. (1867)<br>Brown et al. (2019)<br>Vangelatos et al. (2022) | Diagonal Struts | 1. Improves strength without much redundancy of material.<br>2. Reduce the stresses away from the nodal intersection by delocalizing the stress to neighboring beams. | 1. Mechanical Testing – 3-point bend test, compression test<br>2. Numerical Analysis.<br>3. Simulation.<br>4. Parametric optimization. |
| Weaver et al. (2006)<br>Gray et al. (1867) | Lid and terminal Sleeve | 1. protect the sponge interior and from other aquatic creatures from eating the sponge.<br>2. Prevents ovalization of the sponge.<br>3. Provides stability. | 1. Observed and hypothesized |
| Schulze et al. (1887)<br>Weaver et al. (2006)<br>Gray et al. (1867)<br>Fernandes et al. (2021)<br>Falcucci et al. (2021)<br>Vangelatos et al. (2022) | External Helical Ridge | 1. Resist both failure modes, ovalization and torsion.<br>2. Hydrodynamic benefits - ridge designs are effective at suppressing vortex shedding forcing on the cylindrical structure.<br>3. Used for feeding and sexual reproduction. contact between free-swimming sperm and retained eggs, thereby boosting reproductive efficiency. | 1. Mechanical testing – compression test, torsional test, compression test<br>2. Mechanical and CFD simulations. |

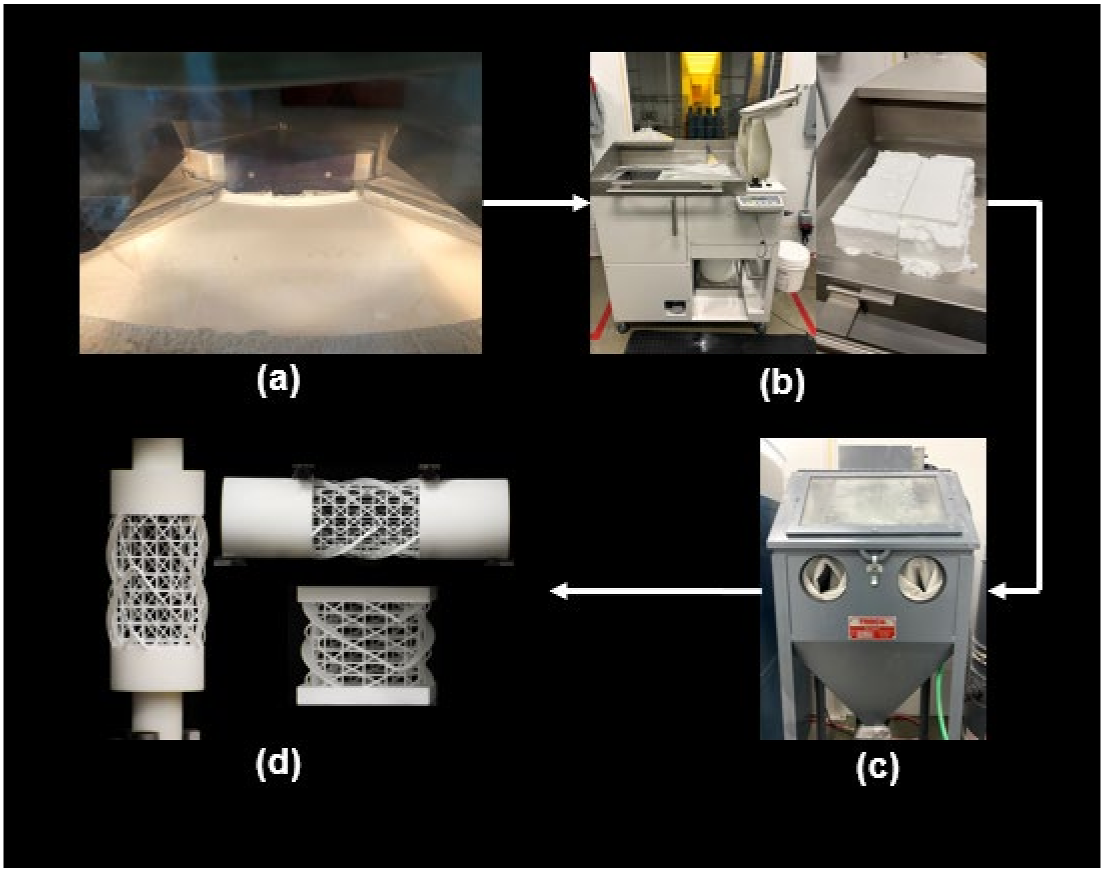


**Figure S1**. 3D printing and post processing of specimens. (a) represents 3D printing of specimens on SLS machine: EOS Formiga P110, (b) represents powder removal and initial cleaning station, (c) represents sand blasting unit, where the trapper powder from the geometry is removed and (d) shows the final specimens ready for testing.
